# Spatio-temporal segmentation of contraction waves in the extra-embryonic membranes of the red flour beetle

**DOI:** 10.1101/2024.08.23.609389

**Authors:** Marc Pereyra, Mariia Golden, Zoë Lange, Artemiy Golden, Frederic Strobl, Ernst H.K. Stelzer, Franziska Matthäus

## Abstract

In this paper, we introduce an image analysis approach for spatio-temporal segmentation, quantification and visualization of movement or contraction patterns in 2D+t or 3D+t microscopy recordings of biological tissues. The imaging pipeline is applied to time lapse images of the embryonic development of the red flour beetle *Tribolium castaneum* recorded with light-sheet fluorescence microscopy (LSFM). We are particularly interested in the dynamics of extra-embryonic membranes, and provide quantitative evidence of the existence of contraction waves during late stages of development. These contraction waves are a novel observation of which neither origin, nor function are yet known. The proposed pipeline relies on particle image velocimetry (PIV) for quantitative movement analysis, surface detection, tissue cartography, and an algorithmic approach to detect characteristic movement dynamics. This approach locates contraction waves in 2D+t and 3D+t reliably and efficiently and allows the automated quantitative analysis, such as the area involved in the contractile behavior, contraction wave duration and frequency, path of contractile area, or the relation to the spatio-temporal velocity distribution. The pipeline will be used in the future to conduct a large-scale characterization and quantification of contraction wave behavior in *Tribolium castaneum* development and can be adapted easily to the identification and segmentation of characteristic tissue dynamics in other systems of interest.

## 1 Introduction

We present a method to detect and quantify spatio-temporal dynamic patterns in biological tissues characterised by bidirectional motion. The method was developed to systematically detect and analyse the recently observed but yet undescribed phenomenon of contraction waves in the extra-embryonic (EE) membranes of the red flour beetle, *Tribolium castaneum*, during its embryonic development.

Contractile cell dynamics are recognized to play important roles during the development (reviewed in: [29,5]) and evolution of species [16,22,1]. In particular, apical contractions of epithelial cells are crucial in morphogenesis [13,14]. Cell contractions have been observed during dorsal closure in the most commonly studied insect model organism, *Drosophila melanogaster*. Here, cells of its single EE membrane, the amnioserosa, [21,3], exhibit apical contractions facilitating a reduction of EE tissue area [4,19,23].

In this work, we study EE membrane dynamics in *Tribolium*. In contrast to *Drosophila*, where the amnioserosa covers only the yolk dorsally, *Tribolium* presents two large and separate EE membranes: the amnion, which covers the embryo itself (embryo proper) ventrally, and the serosa which covers the whole embryo proper and the yolk. The EE membranes eventually rupture in the anteroventral region and rapidly withdraw to the dorsal side. This process signals the onset of dorsal closure [17] which happens roughly one day before hatching. With its architecture and dynamics of EE membranes, *Tribolium* illustrates the embryonic development of the most numerous order of insects, Coleoptera [25]. This makes the insect *Tribolium castaneum* an important emergent model organism for analyzing EE dynamics as exemplified by our novel observation of the propagating contraction waves.

Our image analysis approach consists of three main components that address different aspects of our specific problem. One key feature stems from the geometry of the EE membranes in *Tribolium*. Namely, the serosa EE membrane is a continuous epithelial layer that covers the whole embryo from gastrulation to rupture, which can be geometrically described as a closed surface embedded in 3D space. A powerful approach for analyzing such surfaces is through extracting a surface mask and projecting the intensities onto 2D maps [6]. We included this approach in our approach, as it has the additional advantage of disentangling the dynamics of the EE membranes from movements of the embryo proper.

The second pillar of our approach is ensuring the robust and accurate quantification of tissue dynamics based on microscopy data, as any errors in the quantification would propagate to the wave segmentation algorithm. We use particle image velocimetry (PIV), which is an image analysis technique for computing displacement fields from consecutive images. PIV is commonly used for biological images, since it is accurate at quantifying collective cell motion and can be applied to non-segmentable data [24,2,11]. Both of these merits stem from the fact that PIV uses cross-correlation, which is a translation-invariant pattern-matching operation that produces a maximum peak at the translation that minimises the differences between the cross-correlated images [8]. In particular we use the quickPIV package, which is an efficient and free PIV implementation handling 2D and 3D images. QuickPIV also includes elemental features and options used here, such as masked PIV and normalized cross-correlation using squared differences [18].

Lastly, our approach proposes an original algorithm for segmenting the characteristic back-and-forth motion that we observe during contraction waves in *Tribolium*. We base our algorithm on a set of quantitative angle-based criteria that describe the motion pattern of interest. In particular, we observed from multiple embryos that contraction waves are characterized by alternating dorsal and ventral movements of large patches of EE membranes cells in *Tribolium*. They seem to start in the anterodorsal area and propagate in the posteroventral direction multiple times before the rupture and the withdrawal of EE membranes, which happens at the late stage of embryo development. The algorithm has only a few parameters that intuitively shape the range of detectable patterns. In the following, we introduce the methods that were used to analyse EE membranes dynamics and segment contraction waves. We developed and used methods for both 2D and 3D data and applied them to three different representations of our data: (1) 2D maximum intensity projections of a lateral view of the embryo, (2) 3D fused volumes from time-lapse LSFM 3D recordings, and (3) cylinder projections based on the surface mask where the EE membranes get unrolled and mapped to 2D.

## 2 Methods

### 2.1 Analyzing EE membranes dynamics with PIV

We used quickPIV [18] to quantify tissue motion in 2D and 3D. In the case of the 3D data, we restricted the PIV analyses to the EE membranes by providing a 3D mask of the surface of the embryo. This speeds up the computation and results in PIV vector fields that predominantly capture EE membranes dynamics, and reduce contributions from the movement of the subjacent embryo proper. The 3D surface mask was generated by combining surface masks derived from surface meshes of the embryo at different time points (Section 2.3.3). We used the same mask for all 3D PIV analyses to ensure that all results share the same set of coordinates.

We computed PIV between each pair of consecutive frames for the entire 3D+t microscopy data, as well as 2D+t projections, resulting in 3D+t and 2D+t vector fields, respectively. In the rest of the paper we will refer to an arbitrary vector within the PIV results by its spatial, **p** = (*x, y*) or **p** = (*x, y, z*), and temporal, *t*, coordinates: **v**_**p**,*t*_. In addition, the wave segmentation algorithm revolves around analyzing the change in direction of a PIV vector over time. In other words, the input to the algorithm is the set of PIV vectors at the spatial coordinate **p** across all frames, which we denote by **v**_**p**,{*t*}_. We will refer to this as the “vector time-series at **p**”.

### 2.2 Temporal segmentation of characteristic dynamics based on PIV vector fields

The temporal wave segmentation algorithm was developed based on the *a priori* observation that EE membranes contraction waves in *Tribolium* are defined by two consecutive phases of tissue movement in different directions: an initial phase of posteroventral movement over several frames, followed by a phase of movement with a strong dorsal component for several frames, see Fig. 1A and Fig. 1B. We refer to these two phases as the V and D phases, respectively, highlighting the importance of the ventral and dorsal components of the movements in the contraction waves. This suggests that a time window containing a contraction wave can be identified for a given position by a temporal pattern in the PIV vector at **p** showing the respective characteristic dynamics. To allow for a certain amount of flexibility, as ventral and dorsal movements are not perfectly aligned with the dorsoventral axis, especially after EE membranes rupture, we construct our algorithm around a set of conditions defining the desirable pattern. In particular, we proceed as follows:

**Fig. 1:**
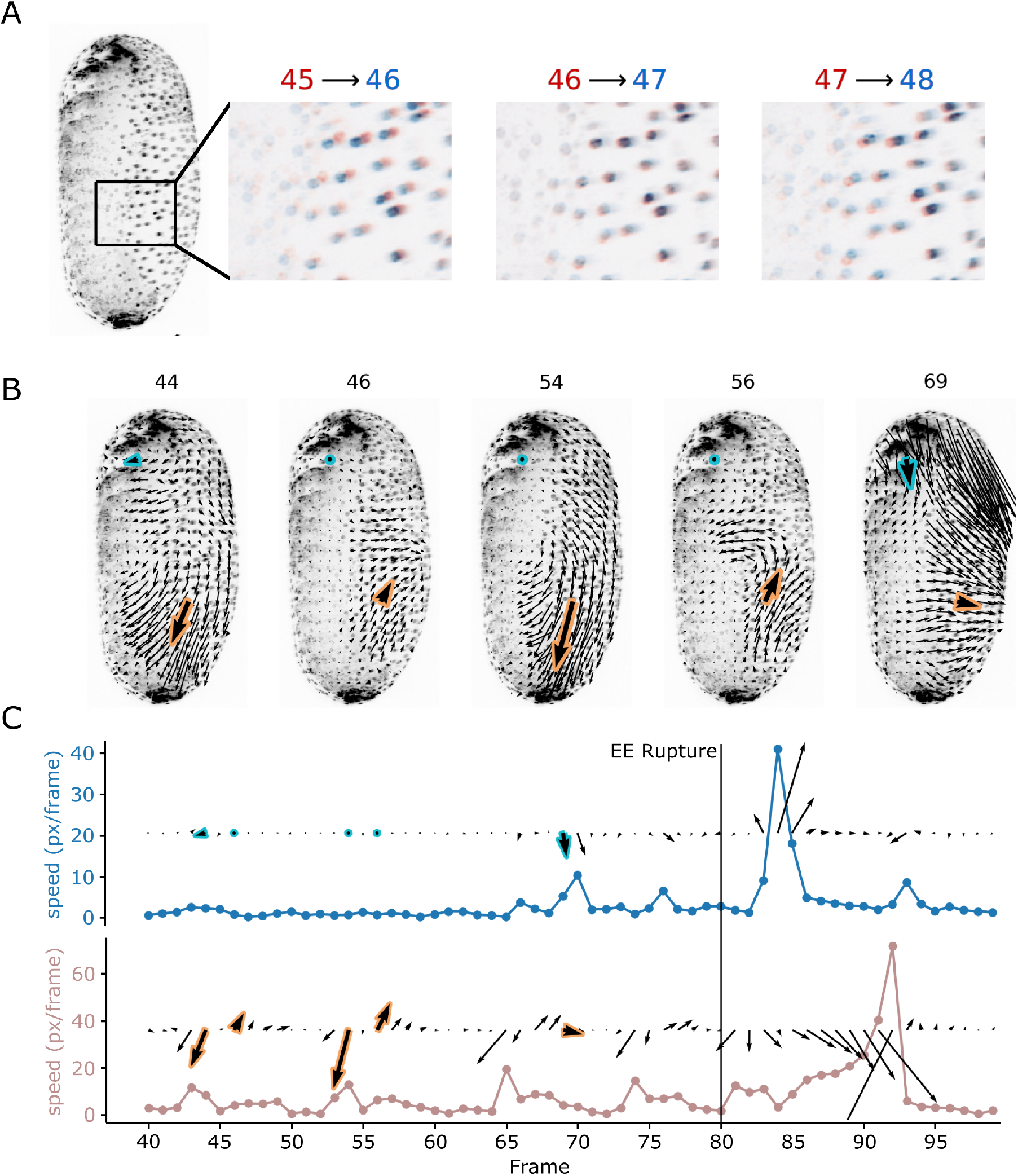
Quantification of contraction waves through PIV. **A** Lateral view of *Tribolium castaneum* (left). Contraction waves are visible from the lateral views as dorsoventral back-and-forth movements of cells of the EE membranen. Each one of the three panels on the right illustrates the movement of the EE membrane cells by overlying the intensities of two consecutive frames: red for the first frame and blue for the second frame. **B** 2D+t PIV results on lateral maximum projections. The contraction waves are captured in the PIV vector fields and are spatially coherent. Two positions of the vector field are highlighted: the blue-bordered vectors, and the orange-bordered vectors. **C** Temporal evolution of the PIV vectors at the highlighted positions in B. The blue vectors do not show any contraction waves, but still show sporadic velocity peaks of about 7-10 px/frame before rupture, as well as a large velocity peak after rupture. The orange vectors illustrate the appearance of PIV vectors in regions of contraction waves. In particular, this vector series shows five waves before rupture and at least one wave after the time of rupture. There is a high velocity peak around frame 90 due to the retracting EE membranes crossing this position of the vector field.

a. We first adopt a sliding window approach to detect frames of maximum and minimum similarity with a reference direction for the V phases.
b. Secondly, we subject each pair of consecutive local maxima and minima from the previous step to a set of angle-based and velocity-based criteria to determine whether they correspond to the V and D phases of a contraction wave. Through this step we obtain temporal segmentations of the waves.
c. After independently generating a temporal wave segmentation for each spatial position of the PIV vector field, we apply post-processing steps to refine the spatial segmentation of the tissue involved in the contraction wave.

#### 2.2.1 Detection of V phase and D phase candidates

The first step in the temporal segmentation of the V-D movement pattern relies purely on direction information. Generally, we are searching in the PIV vector time series **v**_**p**,{*t*}_ for a pattern of *N* consecutive vectors pointing coherently in one direction, followed by *M* vectors pointing coherently in a different direction. In the case of contraction waves in *Tribolium*, the direction of the first phase (the V phase) is constrained to the posterioventral direction. In our 2D lateral projections, we choose the normalized vector representing the angle bisector of the third quadrant (225 degrees, see Fig. 2A). From this reference vector, we define a broader region of allowable directions for the V phases by considering an angular spread of ±40 degrees (*θ*_**r**_ in Fig. 2A). The range of possible directions of the D phase is more variable: it is broadly constrained within 180 degrees of the dorsal direction. In addition, we impose a minimum angle with the average direction of the V phase (*θ*_VD_ in Fig. 2A).

**Fig. 2:**
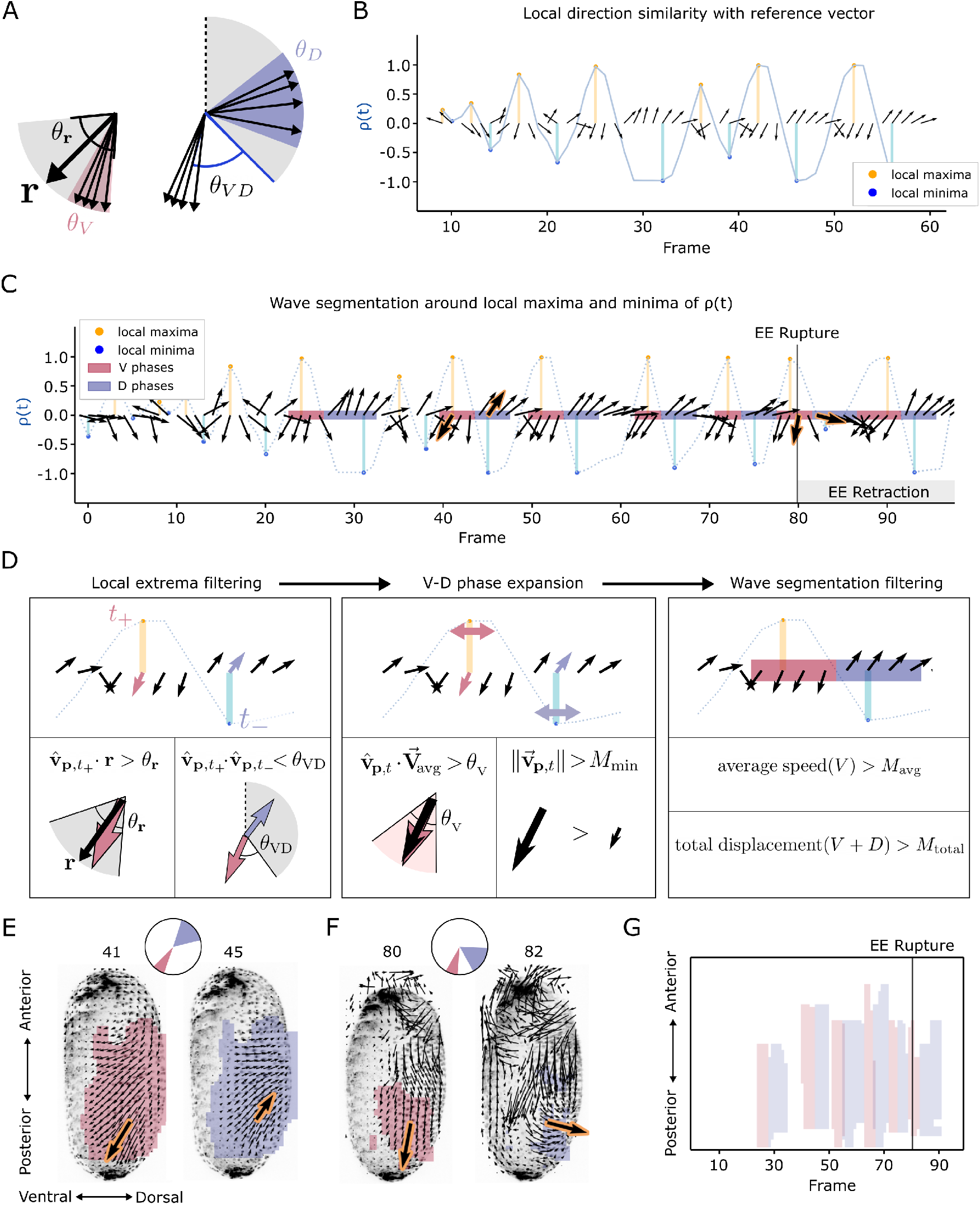
Wave detection algorithm for 2D lateral projections. **A** We define a set of allowable directions for the V phases by considering an angular spread (*θ*_**r**_) around the direction of a reference vector (**r**). The direction of the D phases is restricted by their angle (*θ*_VD_) to the average direction of the preceding V phase. The direction of the D phase is restricted to (−90°, 90°) from the dorsal axis, and by the angle to the average direction of the V phase (*θ*_VD_). We also limit the spread of the vectors within the V and D phases, *θ*_V_ and *θ*_D_.**B** Sliding window quantification of direction similarity (*ρ*(*t*)) on a vector time series, **v**_**p**,{*t*}_, between time points 10 and 60. **C** Wave segmentation results for the same vector time series. **D** The segmentations in panel C were generated starting with pairs of local maxima and minima of *ρ*(*t*) whose directions are consistent with the a-priori angles for the V and D phases (left). The expansions of the V and D segmentations (middle), and their post-processing (left) are parametrized by angle and speed-based criteria. **E** Example of 2D wave segmentation results of a pre-rupture contraction wave. Before rupture, the vectors of the V and D phases point in opposite directions. **F** Example of the 2D wave segmentation of a post-rupture contraction wave. After rupture the angle between the V and D phases is smaller than during pre-rupture waves. **G** Kymograph along the anterior-posterior axes of the wave segmentation results on the 2D+t datasets of maximum projections for all 100 frames.

This pattern can be detected by a sliding window approach on the normalized PIV vector field. For every frame *t* we compute the dot product between the average vector of the normalized PIV vectors around *t*, [*t* − *N, t* + *N*], and the normalized reference vector for the V phase, **r**. Due to the sliding window approach this dot product is again a function of time, denoted as *ρ*(*t*) (Fig. 2B). This measure is maximized if all vectors in the considered window point in the same direction as the reference vector. In other words, *ρ*(*t*) is maximized when the vectors within [*t* − *N, t* + *N*] display low intra-group variability and simultaneously point in the direction of **r**. On the other hand, *ρ*(*t*) is minimized around time points where the PIV vectors in the considered window point collectively away from **r**.

Most notably, *ρ*(*t*) provides an approximate location of the centers of the V and D phases of the waves. The local maxima of *ρ*(*t*) corresponds to time points of coherent movement in a similar direction to **r**, which correlates with the V phases. On the other hand, the local minima of *ρ*(*t*) mostly indicate frames where PIV vectors are coherently pointing in a different direction to *ρ*(*t*), corresponding to the location of the D phases. Local minima can also indicate incoherent movement away from *ρ*(*t*). Thus, we consider each pair of consecutive local maxima and minima as potential landmarks for contraction waves, as illustrated in Fig. 2B.

The normalized reference vector **r** plays a major role in this step and the following step of the wave segmentation algorithm, and it is important to adjust the direction of **r** to each representation of the dataset. In addition, some representations require different reference vectors for different position in the PIV vector field, **r**_**p**_. This is the case for our 3D+t representation and our 2D+t cylinder projections, as the ventral direction at each point is determined by the geometry of each dataset.

#### 2.2.2 Extracting wave segmentations

This step takes each pair of consecutive local maxima and minima (*t*_max_ and *t*_min_) extracted from *ρ*(*t*) and evaluates whether the PIV vectors surrounding these time points fit to the expected pattern of the movement direction characteristic of the contraction waves. If this is the case, a segmentation of the V and D phases is generated. Both of these tasks involve evaluating a set of angle-based criteria: *θ*_**r**_, *θ*_V_, *θ*_D_ and *θ*_VD_. The segmentation step also considers speed-based criteria: *M*_min_, *M*_avg_. Specifically, we evaluate the following set of conditions in the order presented below, where failing any step results in the unsuccessful segmentation of a wave:

1. We first check whether the PIV vector at 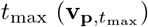 points in the reference direction, **r**. The vector at this frame is characterized by low intra-group angle variances, making it representative of the movement in the V phase. This is tested with a threshold on the normalized dot product: 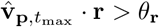 (Fig. 2D left).
2. We then check whether the angle between the representative vectors of the V and D phases is noticeable. The representative vector for the D phase is obtained from 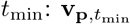. This is tested with a threshold on the following normalized dot product: 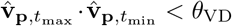 (Fig. 2D left).
3. We proceed by creating a segmentation of the V phase by the process described below (illustrated in Fig. 2D middle panel). The segmentation is initialized to contain a single position, *V* = {*t*_*V*_, }. We expand this segmentation by iterating over the adjacent frames starting at *t*_max_ from nearest to farthest, and deciding whether to add them to *V*. A frame *t* is added to *V* if two conditions are met:
  – the vector at *t* points in a similar direction as the current segmentation of the *V* phase. Since *V* may contain multiple vectors, this is computed with the dot product with the average vector in 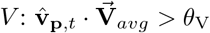
  – the vector’s magnitude is sufficiently large: 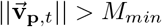
  The same process is applied to generate the segmentation of the D phase, with the only difference of using *θ*_D_ instead of *θ*_V_. If the V and D segmentations are not adjacent, the vectors at the interface are added to the phase with the most similar direction.
4. We impose additional conditions on the segmented *V* and *D* phases. In particular, we discard both the *V* and *D* segmentations if the average speed of the V phase is smaller than a threshold, *M*_avg_. In addition, we include a threshold on the total distance traveled (total displacement) during a wave, which is computed as the sum of magnitudes along the duration of the *V* and *D* phases of a contraction wave.

At the end of the segmentation step, we obtain V and D segmentations for each contraction wave with low intra-angle variance (parameterized by *θ*_V_ and *θ*_D_) and significant inter-angle variance (determined by *θ*_VD_). Speed information is used to avoid extending the segmentation over regions with insufficient movement. The parameters used to segment contraction waves in the different data set representations are provided in Table 1.

**Table 1:** Parameters for wave segmentation algorithm.

|  | Lateral projections | 3D fused volumes | Cylinder projections |
| --- | --- | --- | --- |
| $r_x$ | $-\cos(45^\circ)$ | $-\sin(8.4^\circ)$ | $\pm \cos(45^\circ)$ |
| $r_y$ | $-\sin(45^\circ)$ | $\sin(45^\circ) \cos(8.4^\circ)$ | $-\sin(45^\circ)$ |
| $r_z$ | 0 | $-\cos(45^\circ) \cos(8.4^\circ)$ | 0 |
| $\theta_r$ | $45^\circ$ | $50^\circ$ | $50^\circ$ |
| $\theta_v$ | $60^\circ$ | $60^\circ$ | $40^\circ$ |
| $\theta_D$ | $60^\circ$ | $60^\circ$ | $40^\circ$ |
| $\theta_{VD}$ | $30^\circ$ | $30^\circ$ | $30^\circ$ |
| $M_{\min}$ | 0.5 px/frame | 0.5 vx/frame | 1.0 px/frame |
| $M_{\text{avg}}$ | 1.0 px/frame | 1.0 vx/frame | 3.0 px/frame |
| $M_{\text{total}}$ | 10 pixels | 10 voxels | 10 pixels |

#### 2.2.3 Spatial post-processing of the wave segmentation

The temporal wave segmentation algorithm is independently applied to the vector time-series at each spatial coordinate, **v**_**p**,{*t*}_, and therefore does not include any operations to ensure that wave segmentations are spatially coherent. However, several factors throughout the pipeline impose spatial coherence of the wave segmentation results. Firstly, the phenomenon of interest concerns collective tissue motion, which is inherently spatially coherent. Secondly, spatial coherence is increased as a side product of PIV post-processing filters, such as local spatio-temporal averaging, which is commonly applied to remove high-frequency noise. Therefore, we obtain spatially consistent segmentations during wave events.

However, we also obtained spurious segmentations in other regions of the embryo, either from small wave-like movements in EE membranes or from twitching movements of the embryo proper. We remove small wave-like detections with morphological operations. Namely, we perform morphological opening to the V and D segmentations to remove thin segmentations, followed by a high-pass filter by the size of the connected components in the segmented phases. Artifactual segmentations due to movements in the embryo start after rupture, as EE retraction proceeds and the embryo relocates itself within the egg. We remove these by filtering segmentations whose centroid is near the head of the embryo.

### 2.3 Data processing

#### 2.3.1 Live imaging and fusion of 3D image volumes

Life imaging time lapses of *Tribolium* embryo development were acquired using digitally scanned laser light sheet fluorescence microscopy [9,10]: detection objective 10x 0.3 NA, illumination objective 2.5x 0.06 NA, Andor Clara camera. Mounting and sample preparation was performed as previously described [26], with the addition of fluorescent beads as fiducial markers to aid registrations. The embryo analyzed in this paper was recorded from four directions with rotation steps of 90 degrees at interval of three minutes for a total of 100 frames. For each frame, the four directions were registered and fused into one isotropic volume [7]. This produced a high-quality 3D+t dataset of 100 frames covering the late stages of the development of *Tribolium*.

#### 2.3.2 Lateral maximum intensity projections

In addition to the fused data, we generated a 2D+t dataset consisting of maximum intensity projections from one of the lateral views of the embryo, where the contraction waves are clearly visible.

#### 2.3.3 Surface mesh

Until rupture, which occurs at frame 80, the EE membranes membrane constitute the surface of the embryo. Therefore, we included a data-processing step in our pipeline for extracting the surface of each embryo, allowing us to target our analyses to the EE membranes signal. The surface detection for each volume was implemented as follows: First we generated a binary segmentation of the fused 3D volume into foreground (*Tribolium* signal) and background.

The boundary of this segmentation does not correspond directly to the surface of the embryo, because the foreground segmentation is based on the nuclei signal which is inherently sparse. To obtain a smooth surface, we sampled a 3D point cloud from the segmentation and computed their 3D alpha shape with pymeshlab [15], generating a smooth mesh that connects the segmented nuclei on the surface. Lastly, we ensure to remove facets in the interior of the alpha shape, as these do not correspond to the surface. The surface meshes were used for restricting the PIV analyses to the EE membranes, and also for tissue cartography, which is covered in the next section.

#### 2.3.4 Cylinder projections

To reduce dimensionality and allow analysis with 2D methods, we followed the work of Streichan et al. [6] and used their freely available software “Image Surface Analysis Environment” (ImSAnE). This allowed us to generate an atlas of overlapping maps, in which we could then analyze the surface dynamics of the EE membranes on cylinder projections. Using ImSAnE, we extracted the voxels of interest from the 3D image volume using the surface mesh (Section 2.3.3) and projected the voxel intensities to their respective pixels in a flat 2D geometry. We included 9 radial layers of the mesh, 6 towards the inside and 3 towards the outside, to capture a 10 voxel high radial *z*^*′*^ stack to ensure that we capture all nuclei of EE membranes without including signal from the embryo proper inside. This gave us a stack of 10 2D cylinder maps from which we created a maximum intensity projection. We represented the information on up to two cylinder projections with a 180°rotational offset to each other along the main axis of the mesh. We used the respective maximum intensity cylinder projections to create 2D+t time lapse videos for 2D PIV.

We needed to make a few considerations when computing PIV on cylinder projections, especially when representing the results on 2D maps. In cylinder projections, both distance and angle distortions are introduced. They increase gradually towards the polar region from the equator which has no distortion. We corrected for length distortions introduced by cartography to ensure that elongated or shortened PIV vectors do not impact our wave segmentation. We used a uniform sampling with 10 px distance to create a 2D grid. Using the function *properLength* in ImSAnE we calculated the distance that the sample points would have in 3D space. We obtained two values for each 10×10 tile of the grid: longitudinal distortion (from pole to pole) which is characterized by shortening towards the poles, and latitudinal distortion (parallel to the equator) which is characterized by elongation towards the poles. We then multiplied the distortion fields with the 2D PIV vector field to obtain corrected vector lengths. We do not correct for angle distortions because these only become significant (≫10°) very close to the poles. Instead, we crop the polar regions before analysis.

### 2.4 Visualization

#### 2.4.1 Kymographs

Kymographs are a commonly used method for visualizing spatio-temporal (2D+t) datasets as a 2D plot where one of the axes is the temporal dimension. The other axis of the plot can correspond to a line in the spatial dimension, or a projection of spatial data onto this line. We here used a maximum intensity projection of the data along one of the spatial axes, either X or Y, for the kymograph plots. We use kymographs to show the evolution of the wave segmentation results across time for the 2D+t dataset of maximum projections, Fig. 2, as well as the 2D+t dataset of cylinder projections, Fig. 3. In the case of the maximum projections, we generate kymographs along the anteroposterior axis. This projection illustrates changes over time in the segmented waves along the vertical axis. In the case of the cylinder projections, we show a kymograph along the dorsoventral axis, which depicts the wave segmentations of both sides of the embryos over time.

**Fig. 3:**
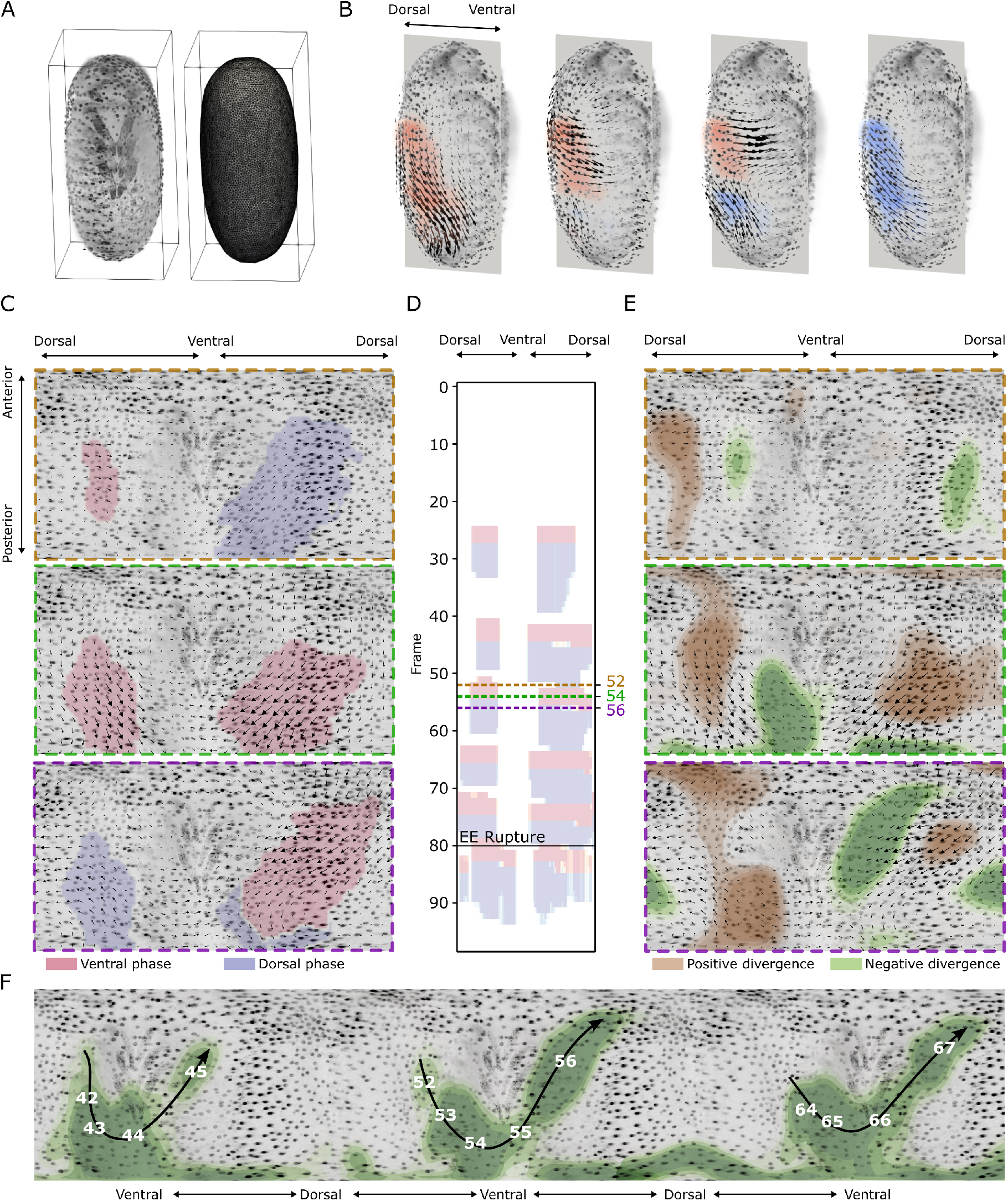
Advanced wave detection pipelines. **A** Example of surface mesh extraction for one frame. We used the detected surfaces both as a mask during 3D PIV analyses and to generate a cartographic representation of the surface. **B** Wave detection on 3D+t vector fields. **C** Wave detection on 2D+t cylinder projections. These projections provide a 360°view of the surface, allowing to detect and visualize contraction waves continuously on both lateral sides of the embryo. The three panels on the left-most column show the progression of one contraction wave. The segmentation on the left side is smaller and precedes the one on the right side. **D** The kymograph summarizes the wave detection for multiple frames, showing the presence of five waves before rupture and one after rupture. **E** In addition, by analyzing the divergence of the PIV vector fields from the cylinder projections, we observed a flow of positive and negative divergence extrema from the left side of the embryo to the right side. This divergence flow correlates with the progression of the wave. **F** Negative divergence (convergence) flows during the three waves prior to rupture. We exploit the periodic boundary of the cylinder projections to visualize the progression of the divergences flow across time by concatenating a projection of frame 54 three times and representing negative divergence calculated on frame 42 up to 67.

#### 2.4.2 Divergence maps

While the focus of this paper is on segmenting the contraction waves, we briefly explored the concurrence between contraction waves and other vector field descriptors. In addition to velocity maps, we computed divergence maps from the PIV result on the 2D cylinder projections. The divergence is useful to identify areas where the tissue density increases (constriction = negative divergence) or decreases (expansion = positive divergence). Divergence is defined as the sum of partial derivatives of each vector field component. This is usually implemented by computing vector field derivatives between adjacent elements. Computing divergence on immediate neighbors requires that the input vector fields be very smooth, which is usually achieved by averaging the vector fields with a kernel of a certain size. In these cases, the size of the averaging kernel implicitly defines the scale of the divergence. Alternatively, we compute divergence by constructing an expanding template vector field of the desired size and cross-correlating this template with each PIV vector field. This expanding template contains normalized vectors pointing away from the center of the template. The size of the template determines the scale of the divergence patterns. We used a kernel size of 17 *×* 17 pixels to compute the divergence on the PIV results generated from the 2D cylinder projections.

### 2.5 Tribolium castaneum husbandry

Live imaging was performed on a transgenic line of *Tribolium castaneum* with fluorescently tagged cell nuclei that expresses histone-binding nanobodies with mRuby attached - ACOS{ATub’H2A/ H2B°NB-mRuby} #1. Beatles cultures were maintained under standard conditions at 32 °C, ∼ 70% relative humidity [27]. For imaging of the contraction waves, beetles were transferred on fresh flour and embryos were collected for 1h at room temperature, then embryos were incubated for 45h at 32. Dechorionation was performed during the second day of incubation.

## 3 Results

### 3.1 Contraction waves correlate with velocity peaks in PIV vector fields

By overlaying the lateral maximum intensity projections of two consecutive frames, we can visualize the dorsoventral shift in individual cell nuclei (Fig. 1A). PIV on the lateral maximum intensity projections captures the contraction waves as large regions of vectors aligned in the dorsoventral axis (Fig. 1B). When observing individual positions of the vector fields within the contraction waves, we find that the pronounced changes in vector direction are accompanied by high velocities. For the temporal resolution used here, the velocities exhibit a peak during the V phase of the waves (shown exemplarily for one position in Fig. 1C). However, also for positions outside the contraction region high velocities are observed, for instance during rapid retraction of the EE membrane after rupture (Fig. 1C). This highlights that velocities alone cannot be used for reliable temporal or spatio-temporal segmentation of the waves.

### 3.2 Analysis of lateral maximum intensity projections yields a segmentation of several contraction waves before rupture and one post-rupture

To segment contraction waves we focus on directional information, as the observed pattern, consisting of a phase of movement in posteroventral direction followed by a phase of movement in dorsal direction, is a more characteristic feature of the contraction waves than local velocity peaks. The proposed algorithm relies on a set of angle-based criteria that define a flexible set of allowable directions for the V and D phases (Fig. 2A). First, we process the PIV vector fields using a sliding window approach (Section 2.2, Fig. 2B). From this, we detect candidate positions for the V and D phases, and then construct the V phase and D phase segmentations iteratively in a second step. In this way, we were able to capture ventral (V) and dorsal (D) phases of variable duration. As a first simple test case, we generated segmentations of the contraction waves both in time (Fig. 2C) and space (Fig. 2 E, F, G) for 2D microscopy data, in particular for a lateral maximum intensity projection of the images.

The chosen set of parameters requires coherence of direction in the V and D phases, but allows for a large flexibility in the angle defining the ventral movement, and also in the relative angles of motion between the V and the D phases. With this, the algorithm can detect pre-rupture waves (Fig. 2E) and post-rupture waves (Fig. 2F) with the same set of parameters, even though the angular spread between the V and D phases in pre- and post-rupture waves is very different. In the kymograph we can identify the presence of five contraction waves before rupture, and one after rupture (Fig. 2G).

After frame 80 the EE membranes rupture and retract to the dorsal side. This involves the collective movement of the entire EE membranes and high velocities at the retracting edge. The retraction phase might also introduce significant ventral and dorsal movement components. However, the choice of parameters (see Table 1) guarantees that contraction waves can be safely distinguished from and are not confused with processes of EE membrane withdrawal.

### 3.3 Wave segmentation on the full EE surface illuminates the spatial characteristics of the contraction wave

In the next step we performed an analysis on the full spatial dynamics of the contraction waves in 3D. Because we were only interested in the EE membrane, we extracted the surface from fused 3D image data (Section 2.3.3, Fig. 3A) and used it as a mask for PIV analysis. Masked data has multiple benefits: (1) the input data for 3D PIV is smaller, making PIV computationally cheaper and faster, and (2) the mask reduces interfering signals, e.g., motion of embryo proper, which might impact PIV results due to signal entanglement.

We performed 3D PIV on the masked 3D image, and used the same parameters for contraction wave detection and segmentation as for the lateral projections (Section 2.3.2). In particular, we used a single static reference vector **r** pointing in ventral direction. With this approach the contraction waves are detected, and can be segmented and visualized also in 3D (see Fig. 3B). PIV analysis and subsequent contraction wave segmentation in 3D showed that the contraction waves occur on both lateral sides and are largely synchronized. With the static reference vector, however, we expect to capture only a fraction of the tissue area involved in the contraction dynamics as the movement occurs on a curved surface. Hence, movement towards the ventral area involves a dorsal-ventral direction only in lateral regions, while in ventral and dorsal areas lateral components dominate. A possible solution to better capture the 3D dynamics would be to use a spatially resolved reference vector field tangential to the surface.

Instead, we decided to adopt tissue cartography, a method to represent data from a (closed) surface embedded in 3D by projection onto 2D maps (Fig. 3C). PIV was performed on the 2D maps, using correction of the length distortions (Section 2.3.4). For wave segmentation we used two reference vectors, **r**_1_ and **r**_2_, for the vectors on the right and left side of the embryo. The direct visualization of the contraction waves on the mapped surface, as well as a kymograph along the dorsoventral axis allow for a comparison of the waves on the left and the right side of the embryo (Fig. 3D). In this dataset, the contraction waves on the left side precede the waves on the right side (Fig. 3D). This phase shift was very small (one frame, i.e., 180 s). The tissue area involved in the contraction waves on both lateral side also differs, with the area on the left being consistently smaller than the segmented area on the right (Fig. 3C). In addition, the kymograph (Fig. 3D) shows that the duration of the V phase is generally shorter than the D phase, a dynamics indicating an abrupt contraction followed by an extended relaxation phase. This consistent signature of the two contraction wave phases is also supported by the velocity dynamics at individual positions, as shown in (Fig. 1C).

### 3.4 Repetitive flow pattern of divergence minima coincides with the progression of contraction waves

Finally, we used the divergence of the vector field to identify tissue regions in the EE membranes that, by local constriction (negative divergence) and expansion (positive divergence), could potentially generate the observed waves. We quantified the local divergence minima and maxima during the frames of ongoing contraction waves and overlaid them on the cartography maps (Fig. 3E). Coinciding with the small phase shift in the onset of the ventral phase of the contraction waves between the two sides of the embryo (Section 3.3), we found a characteristic pattern in the spatial localization of the divergence minima for several consecutive time frames (Fig. 3E). Specifically, the minima moved from the left side of the embryo proper, down to the posterior pole, and up to the right side, coinciding with the progression of a contraction wave and being repeated for each detected wave in a similar manner, as shown by augmenting cylinder projections (Fig. 3F).

## 4 Conclusion

We present an approach for the spatio-temporal analysis of movement patterns in 3D biological tissues that involves movement quantification by PIV, segmentation of tissue areas showing characteristic movement behavior, and different representations of 3D microscopy data. Here, we applied it to detect contraction waves in the extra-embryonic membranes of the beetle *Tribolium castaneum*. Central to our approach is the algorithm for the detection and segmentation of biphasic movements in 2D+t and 3D+t biological datasets.

This algorithm is based on selecting a flexible set of rules to detect *a priori* defined patterns. These rules combine angle-based and speed-based criteria, since velocities alone are not enough to differentiate between EE contraction waves and other rapid developmental events, e.g. EE membranes rupture and retraction. The algorithm includes only a few parameters that control the tolerances, which allowed us to reliably detect both pre- and post-rupture contraction waves. The fact that the algorithm accepts a separate reference vector for each position of the vector field opens interesting possibilities in the future, e.g. the possibility to define multiple locations around which contraction waves may be detected.

We applied the wave segmentation approach separately to three different representations of our *Tribolium* dataset: (1) lateral maximum intensity projections, (2) surface voxels of 3D fused data, (3) cylinder maximum intensity projections. For each of these representations we defined a set of reference vectors indicating the wave direction we want to segment. The lateral maximum intensity projections require only one reference vector, allowing us to quantify a change in the contraction wave orientation after EE membranes rupture. To study the full spatial characteristics of the contraction wave, we moved to fused 3D data. In a 3D representation of the data, however, we found that using only one reference vector is not sufficient to segment the full area of a wave due to Gaussian curvature. Here, each vector field position would require a unique reference vector tangential to the surface. Instead of expanding to a field of reference vectors, we created cylinder projections on which we could define two mirrored reference vectors and corrected our wave segmentation for distortions [6]. The advantage of this approach is that we unroll the EE membranes into a flat 360 degree longitudinal representation allowing for a curvature-free 2D analysis of the surface around the equator and removing potentially coinciding signal from the bulk, specifically the embryo proper.

Additionally, cylinder projections enabled us to track a characteristic “U”-shaped flow of negative divergence in consecutive frames, which was characteristic of multiple waves: the contraction propagates posteriorly over one side of the embryo, crosses the ventral line, and moves anteriorly over the other side. This might be indicative of a propagating focus of contractility that tugs on the neighboring tissue and causes the displacement of nuclei, which we perceive as a contraction wave. The trajectory of this contractile focus is non-intuitive, and might hint at a more complex interaction between geometry, structural and regulatory elements involved in the contraction wave dynamics in this organism. In any case, the repetitive unidirectional shift of the divergence minima may be an important biological insight and needs to be supported on a bigger sample size.

The method presented here in combination with high throughput imaging techniques and 2D/3D PIV [18] will be used in our future work to not only qualitatively characterize and segment the contraction waves but to statistically evaluate the count, size and shape of contraction waves over multiple waves in multiple samples. With this, we hope to answer questions revolving around contraction wave dynamics: e.g. How many waves occur before and after rupture? Does the area of the waves differ between consecutive wave cycles? Do they change their shape, orientation or location? The computing time and robustness of the spatio-temporal segmentation approach make it possible to apply it to multiple 3D datasets, acquired with, e.g., light-sheet fluorescence microscopy that produces image volumes of 1-5 Terabyte per dataset. This makes the method also relevant for smart microscopy where a faster acquisition mode is triggered by a characteristic spatio-temporal pattern [12], and therefore allows to image rapid processes of EE membranes, such as rupture and retraction, with increased temporal resolution. Moreover, the 3D version of this method does not necessarily have to be used on the surface of the sample. Instead, the surface layer can be “peeled off” to allow 3D analysis of the bulk of the embryo in order to potentially identify wave-like dynamics of the embryo proper or the yolk.

Similar waves have been studied in other distant biological systems, e.g., a simple marine animal *Trichoplax adhaerens* [1] and an agglomeration of sea star embryos that form a confluent sheet [28], where waves occur in the presence of global movements, i.e., locomotion and cluster rotation, that could be disentangled by moving to a relative reference frame such that the relative (angular) velocity is zero. Our algorithm, however, was designed to segment contraction waves in the presence of local spatio-temporal dynamics that are prevalent in multi-layered embryonic systems, such as the movement of the yolk and the embryo proper underneath the EE membranes, which required an approach based on angles and velocities. In general, the wave detection presented in this work, is applicable to other multi-cellular or multi-agent systems and is extendable for other spatio-temporal patterns in case they are collective and quantifiable by PIV, e.g., rhythmic yolk contractions in goldfish eggs and embryos [20].

## Notes

### Competing Interest Statement

The authors have declared no competing interest.

